# Dietary restriction of isoleucine increases healthspan and lifespan of genetically heterogeneous mice

**DOI:** 10.1101/2022.10.06.511051

**Authors:** Cara L. Green, Michaela E. Trautman, Reji Babygirija, Raghav Jain, Krittisak Chaiyakul, Heidi H. Pak, Anneliese Bleicher, Grace Novak, Michelle M. Sonsalla, Chung-Yang Yeh, Mariah F. Calubag, Teresa T. Liu, Victoria Flores, Sarah Newman, Will A. Ricke, Kristina A. Matkowskyj, Irene M. Ong, Judith Simcox, Dudley W. Lamming

**Affiliations:** Department of Medicine, University of Wisconsin-Madison, Madison, WI 53705, USA; William S. Middleton Memorial Veterans Hospital, Madison, WI 53705, USA; Interdepartmental Graduate Program in Nutritional Sciences, University of Wisconsin-Madison, Madison, WI 53706, USA; Graduate Program in Cellular and Molecular Biology, University of Wisconsin-Madison, Madison, WI 53706, USA; Department of Biochemistry, University of Wisconsin-Madison, Madison, WI 53706, USA; Integrated Program in Biochemistry, University of Wisconsin-Madison, Madison, WI 53706, USA; Department of Biostatistics and Medical Informatics, University of Wisconsin-Madison, WI 53705, USA; Comparative Biomedical Sciences Graduate Program, University of Wisconsin-Madison, Madison, WI 53706, USA; George M. O’Brien Center of Research Excellence, Department of Urology, University of Wisconsin, Madison, WI, 93705, USA; Department of Pathology and Laboratory Medicine, University of Wisconsin-Madison, Madison, WI, USA; University of Wisconsin Carbone Comprehensive Cancer Center, University of Wisconsin, Madison, WI 53705, USA; Department of Obstetrics and Gynecology, University of Wisconsin-Madison, Madison, WI 53705, USA

**Keywords:** isoleucine, lifespan, frailty, mice, branched-chain amino acids

## Abstract

Low protein (LP) diets promote health and longevity in diverse species. Although the precise components of an LP diet that mediate its beneficial effects have not been defined, reducing dietary levels of the three branched-chain amino acids (BCAAs) leucine, isoleucine and valine promotes metabolic health in both sexes, and increases lifespan while reducing frailty in male, but not female, C57BL/6J mice. Each BCAA has unique metabolic effects, and we recently showed that restriction of isoleucine is both sufficient to promote metabolic health and required for the metabolic benefits of an LP diet in male C57BL/6J mice. Here, we tested the hypothesis that specifically restricting isoleucine could promote healthy aging in genetically heterogenous UM-HET3 mice. We find that a reduced isoleucine diet improves the metabolic health of both young and old HET3 mice, promoting leanness and glycemic control. Restriction of isoleucine starting in adult, 6 month old HET3 mice reprograms hepatic metabolism in a way distinct from an LP diet. Finally, we find that a reduced isoleucine diet reduces frailty and extends the lifespan of both male and female HET3 mice, but to a much greater degree in males. Our results demonstrate that restricting dietary isoleucine can increase health span and longevity in a genetically diverse population of mice, and suggests that reducing dietary levels of isoleucine may have great potential as a geroprotective intervention.

## Introduction

Dietary interventions can promote healthy aging and even extend lifespan. The most well-characterized of these interventions is calorie restriction (CR), which extends the lifespan and healthspan of diverse species, including rodents and non-human primates (Colman et al., 2014; McCay et al., 1935; Osborne et al., 1917; Weindruch et al., 1986). As a CR diet may be too difficult for most people to follow, there has been a great deal of interest in developing interventions that can mimic the benefits of a CR diet without requiring reduced calorie intake.

Dietary composition has a strong influence on longevity and health. Multiple short term randomized clinical trials of protein restriction (PR) show that PR improves multiple markers of metabolic health, reducing adiposity and improved insulin sensitivity (Ferraz-Bannitz et al., 2022; Fontana et al., 2016). In rodents, PR improves metabolic health, and the consumption of low protein (LP) diets promotes longevity (Green and Lamming, 2019; Green et al., 2022; Laeger et al., 2014; Richardson et al., 2021; Solon-Biet et al., 2014; Speakman et al., 2016).

While the precise mechanisms which mediate the beneficial effects of CR and PR have not been defined, these diets inevitably involve reduced consumption of essential dietary amino acids. Restriction of the nine essential amino acids is required for the benefits of a CR diet on lifespan (Yoshida et al., 2018), suggesting that one or more of the essential amino acids can regulate lifespan. While significant attention has focused on methionine, restriction of which can extend the lifespan of flies and rodents (Lee et al., 2014; Miller et al., 2005; Orentreich et al., 1993), we have shown that in the context of an LP diet, the branched-chain amino acids (BCAAs) leucine, isoleucine, and valine are of unique metabolic importance. Restriction of the three BCAAs by 67% - from the level found in a Control diet where 21% of calories are derived from protein to the level found in a LP diet where 7% of calories are derived from protein – improves metabolic health in both sexes of mice, and in male C57BL/6J mice extends lifespan by over 30%, equivalently to an LP diet (Richardson *et al*., 2021).

While the three BCAAs have historically been considered as a group, it is now clear that each of the three BCAAs have distinct molecular and metabolic effects; this may be due to differential sensing of the BCAAs, for example by the mTOR protein kinase (Wolfson et al., 2016), or due to the fact that the intermediate and final products of each BCAA are distinct. For example, 3-hydroxy-isobutyrate is a valine-specific catabolite that regulates trans-endothelial fatty acid transport and glucose uptake (Bishop et al., 2022; Jang et al., 2016). We recently showed that the beneficial metabolic effects of protein or BCAA restriction are principally mediated by restriction of isoleucine. Restriction of isoleucine alone promotes glucose tolerance, improving hepatic insulin sensitivity, and reduces adiposity (Yu et al., 2021). Importantly, restriction of isoleucine is necessary for the metabolic benefits of an LP diet (Yu *et al*., 2021). Isoleucine may also play a critical role in human health and aging; dietary isoleucine levels are associated with human body mass index, while blood levels of isoleucine correlate with increased mortality (Deelen et al., 2019; Yu *et al*., 2021).

Here, we investigated the hypothesis that reduced isoleucine consumption could extend the healthspan and lifespan of mice. We conducted this study using UM-HET3 (HET3) mice, a defined and genetically heterogeneous background which has been extensively used in aging studies (Flurkey et al., 2010; Harrison et al., 2009; Miller et al., 2011; Strong et al., 2016). In addition to better reflecting the genetically diverse human population than any single inbred strain, using HET3 mice increases the likelihood that any salient findings will be generalizable and robust, and not related to a strain-specific genetic defect or cause of death. We find that short term restriction of isoleucine by 67% improves metabolic health in young HET3 mice of both sexes, reducing weight, adiposity, and improving glucose homeostasis. Long-term dietary restriction of isoleucine in adult HET3 mice promotes lifelong leanness and glycemic control, and reprograms hepatic metabolism in a way distinct from a diet in which all dietary amino acids have been restricted. Finally, we show that lifelong restriction of isoleucine by 67% starting at 6 months of age, but not a diet in which all amino acids are restricted to the same degree, reduces frailty and extends median and maximum lifespan in male mice, and median lifespan in female mice, although to a greater degree in males than in females. In conclusion, our results demonstrate that reducing dietary levels of isoleucine promotes metabolic health, reduces frailty and increases longevity in mice, is more robust than a low protein diet in its ability to promote healthy aging, and may be a uniquely potent method for preventing and intervening in age-related disease without requiring reduced calorie consumption.

## Results

### Isoleucine restriction reduces body weight and adiposity and improves glycemic control in young HET3 mice

We first placed male and female 9-week-old HET3 mice (HET3 mice are the F2 progeny of (BALB/cJ × C57BL/6J) mothers and (C3H/HeJ × DBA/2J) fathers, and have segregating alleles from all four parental strains) on one of three amino acid (AA)-defined diets that we have previously published (Yu *et al*., 2021). Briefly, our Control diet contains all twenty common AAs; the diet composition reflects that of a natural chow diet in which 21% of calories are derived from protein. We also utilized diets in which all AAs (Low AA) or specifically isoleucine (Low Ile) was reduced by 67%. All three of these diets are isocaloric, with identical levels of fat; in the case of the Low AA diet, carbohydrates were used to replace calories from AAs, while in the case of the Low Ile diet, non-essential AAs were increased to keep the calories derived from AAs constant (**Fig. 1A, Table S1**).

**Figure 1:**
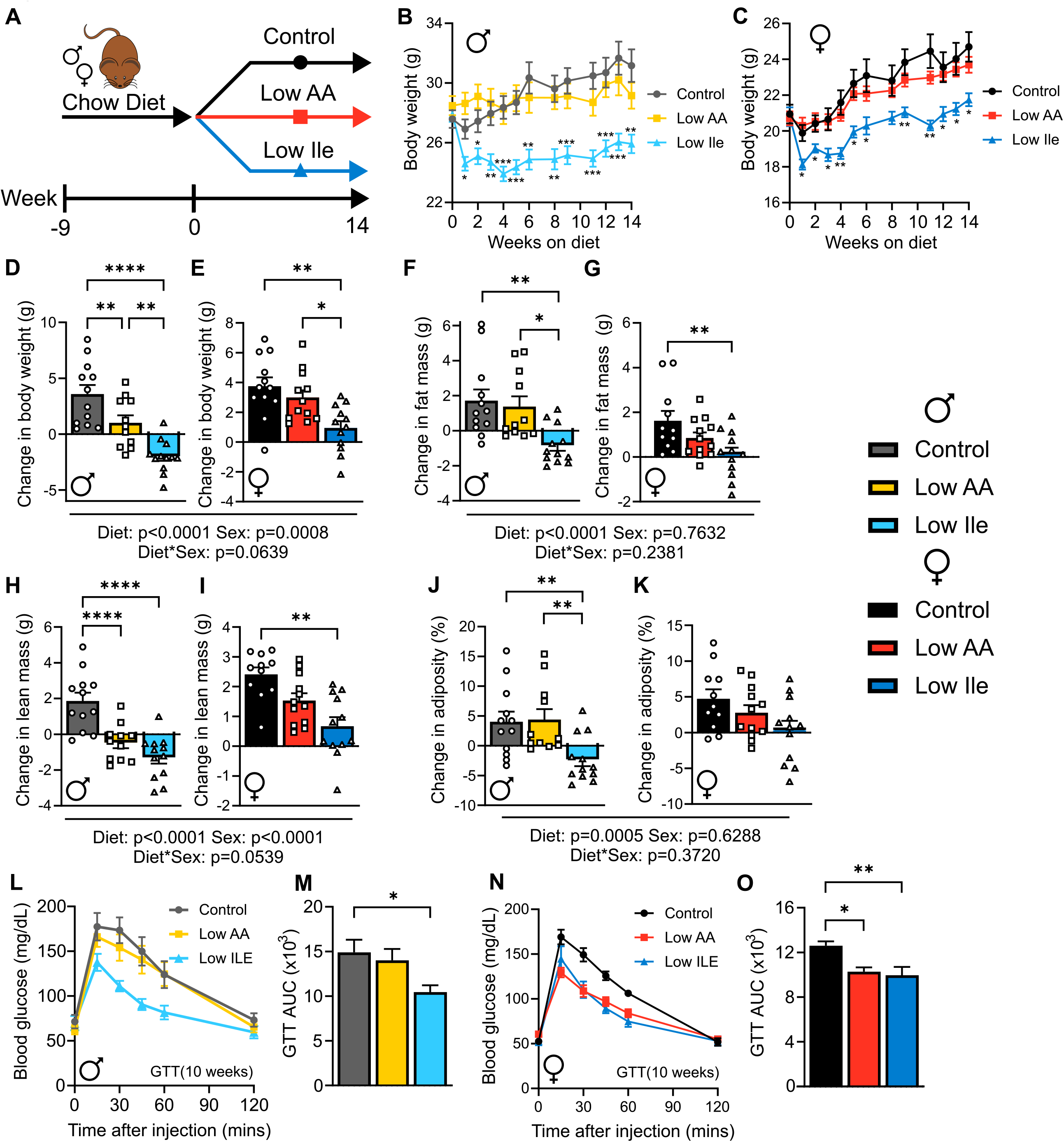
A Low Ile diet reduces body weight and improves glycemic control in male and female HET3 mice. (A) Experimental plan for short-term Low Ile study; male and female HET3 mice were fed the indicated diets starting at 9 weeks of age. (B) Male mouse weight over the course of the 14-week experiment (C) Female mouse weight over the course of the 14-week experiment. (D-K) For males and females respectively, the change in weight (D-E), fat mass (F-G) lean mass (H-I) and adiposity (J-K) over the course of the experiment. (L) Male glucose tolerance test (GTT) after 10 weeks on diet and (M) representative area under the curve (AUC). (N) Female glucose tolerance test (GTT) after 10 weeks on diet and (O) representative area under the curve (AUC). (B-O) n=11-12 mice/group. (B-C) Mixed-effects model (REML) for time and diet with post-hoc Tukey test,\**P*<0.05, \*\**P*<0.01, \*\*\**P*<0.001. (D-K) Two-way ANOVA between Sex and Diet groups with post-hoc Tukey test for pairwise comparisons. (M,O) One-way ANOVA for effect of diet with post-hoc Tukey test for pairwise comparisons, \**P*<0.05, \*\**P*<0.01, \*\*\**P*<0.001 and \*\*\*\**P*<0.0001. P-values for the overall effect of Sex, Diet and the interaction represent the significant p-values from the two-way ANOVA. Data are represented as mean ± SEM.

Over 14 weeks on diet, both male (**Fig. 1B**) and female (**Fig. 1C**) mice on Control and Low AA diets gained weight; however Low Ile-fed mice of both sexes lost a significant amount of weight during the first week on diet (∼12% in males and ∼9% in females), which was maintained across the following 3 months, with female mice gradually gaining some of the weight back (mixed-effects model (REML) for diet, time and the interaction, all significant (p<0.05) for both male and female mice). Over the course of 24 months, the change of weight was time and diet-dependent; male and female mice fed the Low Ile diets, as well as male mice fed a Low AA diet, gained significantly less weight than their Control-fed counterparts (**Figs. 1D-E)**.

Body composition was determined at the beginning and end of the experiment. We found a significant effect of Low Ile on fat mass and lean mass (**Figs. 1F-I**), which was diet dependent, and sex dependent in terms of lean mass only (two-way ANOVA for diet and sex). Low Ile-fed males accreted significantly less fat mass and lean mass during the course of the experiment, actually losing fat mass, while Control and Low AA-fed males gained fat mass (**Fig. 1F, H**). Female Low Ile mice also accreted significantly less fat mass than Control and Low AA mice (**Fig. 1G**). Male Low AA and Low Ile-fed mice lost lean mass, while Low Ile-fed female mice accreted significantly less lean mass (**Fig. 1H-I**). Importantly, the overall effect of the Low Ile diet was reduced adiposity, which reached statistical significance in male mice; intriguingly, Low AA-fed mice were not significantly leaner than Control-fed mice (**Figs. 1J-K and S1A-B**).

We expected that a Low Ile and a Low AA diet would improve glycemic control, and performed a glucose tolerance test (GTT) after 3 weeks (**Figs. S1C-D**) and 10 weeks on diet (**Figs. 1L-O**). The effects of the diets on glucose tolerance were dependent on both diet and sex (two-way ANOVA for diet and sex), with males on Low Ile showing significant improvements at both time points whereas females did not. This was independent of improvements in insulin sensitivity, as measured through an insulin tolerance test (ITT) after either 4 weeks or 11 weeks on diet (**Figs. S1E-H**), although female mice on the Low Ile diet had improved insulin sensitivity at 4 weeks relative to Low AA-fed mice. Looking specifically at hepatic gluconeogenesis through intraperitoneal administration of L-alanine in alanine tolerance tests (ATTs), showed that short term Low Ile feeding appeared to improve hepatic gluconeogenesis in both male and female mice, but this effect was temporal, as significance had disappeared after 12 weeks on diet (**Figs. S1I-L**).

### Isoleucine restriction increases food intake and energy expenditure in young HET3 mice

Despite the effects of a Low Ile diet on body weight and composition, Low-Ile fed males (**Fig. 2A**) and females (**Fig. 2B**) ate significantly more calories each day than Control-fed mice, as did Low AA fed mice of both sexes. As a result, both male and female mice fed the Low Ile diet had significantly greater total intake of amino acids than Control-fed mice (**Figs. S2A-B**), while still consuming significantly less isoleucine (**Figs. 2C-D**). Notably, Low AA-fed mice of both sexes consumed a similar amount isoleucine as Low Ile-fed mice.

**Figure 2:**
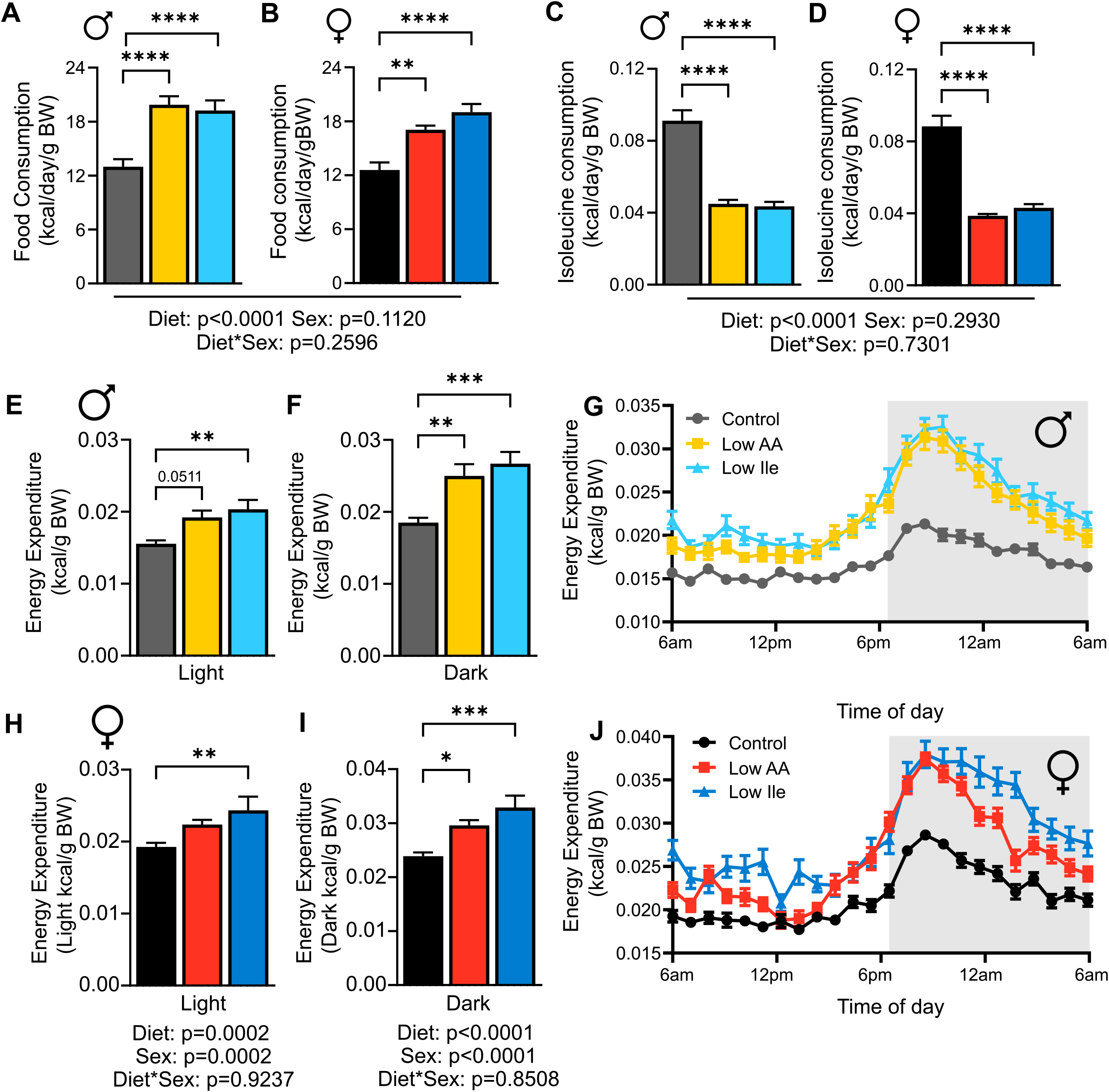
A Low Ile diet increases food consumption and energy expenditure in male and female HET3 mice. (A) Food consumption was measured in home cages after 3 weeks on diet in male mice. (B) Food consumption was measured in home cages after 3 weeks on diet in female mice. (C) Calculated isoleucine intake for male mice. (D) Calculated isoleucine intake for female mice. (E-F) Respectively energy expenditure measure in the light and dark phases in male mice. (G) Energy expenditure of a 24 hour period in male mice. (H-I) Energy expenditure measured respectively in the light and dark phases in female mice. (J) Energy expenditure of a 24-hour period in female mice. Two-way ANOVA for Diet and Sex with post-hoc Tukey multiple comparison test, \**P*<0.05, \*\**P*<0.01, \*\*\**P*<0.001 and \*\*\*\**P*<0.0001. P-values for the overall effect of Diet, Sex and the interaction represent the significant p-values from the two-way ANOVA. Data are represented as mean ± SEM.

Low Ile-fed mice weigh less than Control-fed mice despite significantly increased calorie intake, and we therefore investigated other components of energy balance using metabolic chambers. We found that male mice on a Low Ile diet had drastically increased energy expenditure during both the light and dark cycles; the effect was particularly prominent during the “active” dark phase, where Low AA fed mice also had significantly upregulated energy expenditure (**Figs. 2E-G**). Female mice displayed a similar pattern, with increased energy expenditure particulate during the dark phase in Low Ile and Low AA-fed mice (**Figs. 2H-I**). We did not see any changes in respiratory exchange ratio (RER) in either the light or dark phase (**Figs. S2C-D**); however, there was a shift in the transition from low to high RER, with Low Ile-fed mice shifting from low RER to high RER several hours later than Control or Low AA-fed mice (**Figs. S2E-F**). The alterations in feeding activity and energy expenditure was not associated with changes in activity levels (**Figs. S2G-J**).

### Isoleucine restriction reduces body weight and adiposity and improves glycemic control in old HET3 mice

We next sought to identify if a Low Ile diet can improve healthspan and extend lifespan starting in adult mice. Male and female HET3 mice raised on a chow diet were randomized to three groups of equal weight for each sex, and then placed on either Control, Low AA, or Low Ile diets at 6 months until 24 months of age; at 24 months of age, a pre-selected group of animals was euthanized, and tissues collected, while the remaining mice were allowed to continuing aging (**Fig. 3A**). Tracking weight longitudinally, we found that Low Ile-fed male and female mice both weighed significantly less than both the Control AA and Low AA-fed mice (**Figs. 3B-C**).We observed that much of the difference in male mice was due to a rapid loss of fat mass (∼50% in Low Ile-fed mice) during the first month, while much of the difference in female mice was due to a gain of fat mass in the Control-fed animals (**Figs. 3D-E**). Differences in lean mass contributed much less to the difference in weight in both sexes, with male Low Ile fed mice having an ∼10% drop in lean mass during the first month which was maintained until late life (**Figs. 3F-G**).

**Figure 3:**
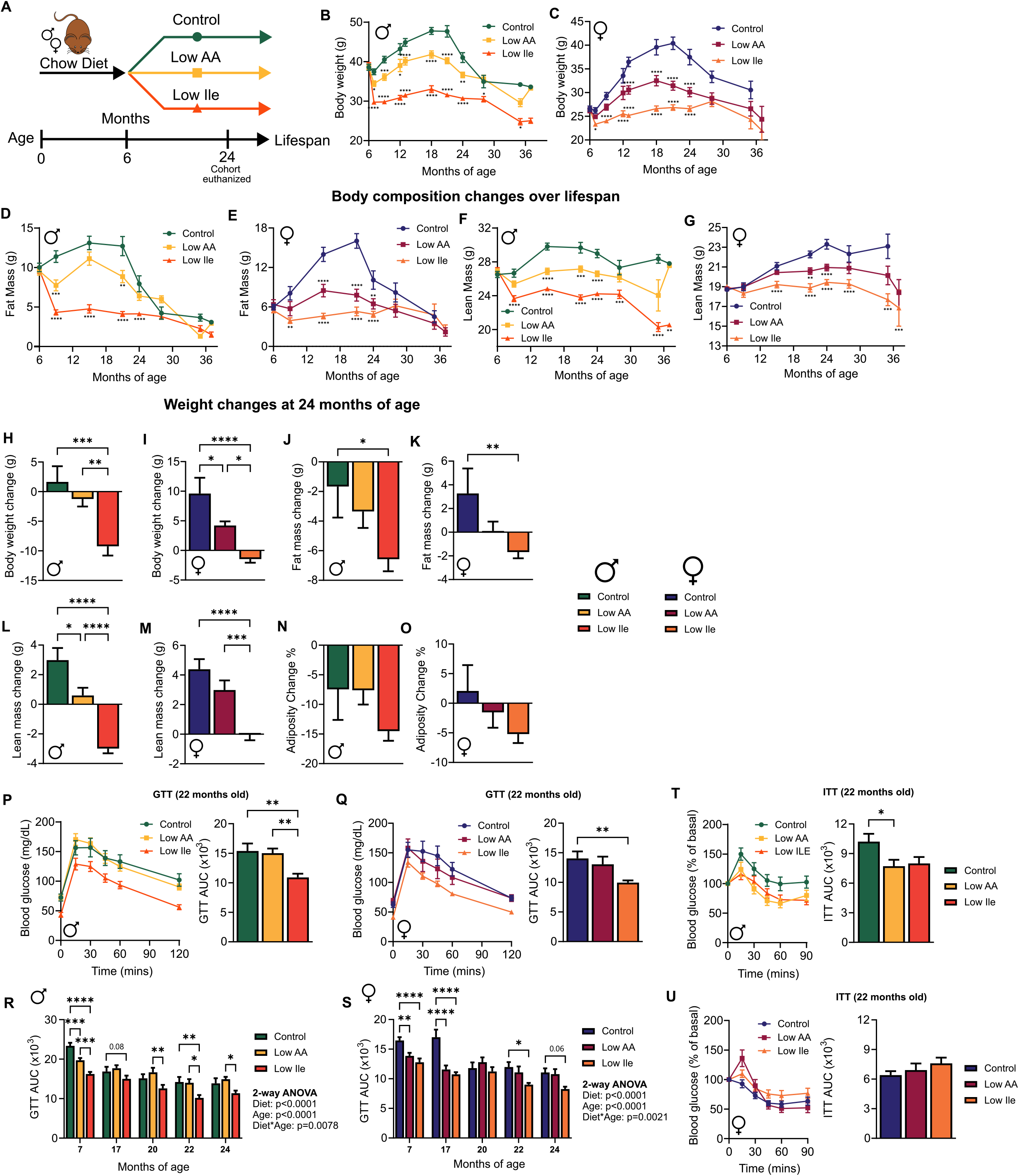
A Low Ile diet reduces body weight and improves glycemic control in male and female HET3 mice when started in mid-life. (A) Experimental plan for long-term Low Ile study; male and female HET3 mice were fed the indicated diets starting at 6 months of age. At 24 months of age, a pre-selected cohort of mice in each group was euthanized for detailed analysis. (B-C) Male (B) and female (C) weight was tracked over the course of the lifespan. (D-G) Body composition was determined over the course of the lifespan with fat mass (D-E) and lean mass (F-G) determined in both male and female mice as indicated. (H-O) For males and females sacrificed at 24 months as indicated the change in weight (H-I), fat mass (J-K) lean mass (L-M) and adiposity (N-O) between 6 months (diet start) and 24 months of age. (P-Q) Glucose tolerance test (GTT) after 16 months (22 months of age) on the indicated diet in male (P) and female (Q) mice with area under the curve at that age (AUC). (R-S) GTT AUC for male and female mice over the course of the experiment. (T-U) Insulin tolerance test (ITT) after 16 months on diet (22 months of age) and representative AUC. n=47-53 mice/group at beginning of study; exact n varies across time points. (H-Q,T-U) One-way ANOVA for diet with post-hoc Tukey test, \**P*<0.05, \*\**P*<0.01, \*\*\**P*<0.001 and \*\*\*\**P*<0.0001. (R-S) Two-way ANOVA for Age and Diet with post-hoc Tukey test, \**P*<0.05, \*\**P*<0.01, \*\*\**P*<0.001 and \*\*\*\**P*<0.0001. P-values for the overall effect of Diet, Age and the interaction represent the significant p-values from the two-way ANOVA. Data are represented as mean ± SEM.

Between 20-26 months of age, we see frailty related declines in body weight and composition, therefore we looked more closely at changes at 24 months of age, prior to euthanizing an experimental cohort of mice. We found that Low Ile-fed male and female mice had an overall reduction in weight during this period, while Control-fed animals overall gained weight (**Figs. 3H-I**). In both sexes, this was due principally to a difference in fat mass accretion, with Low-Ile fed males losing a large amount of fat and Low Ile-fed female mice not gaining fat mass while Control-fed females did (**Figs. 3J-K**). Male, but not female, Low Ile-fed mice also lost lean mass relative to baseline (**Figs. 3L-M**). There were no significant differences in adiposity between diet groups in male mice (**Fig. 3N**), which all lost adiposity between 6 and 24 months of age, or between female mice on (**Fig. 3O**).

We performed glucose and insulin tolerance tests at multiple time points during the lifespan, up to 24 months of age. At 22 months of age (after 16 months on the indicated diets), we found that a Low Ile diet, but not a Low AA diet improved glucose tolerance in both male and female mice (**Figs. 3P-Q**). Looking over time, we find that both Low AA and Low Ile diets improved glucose tolerance earlier in life in both male and female mice, but that this effect as persistent only in Low Ile-fed mice (**Figs. 3R-S**). Interestingly, Low AA-fed male mice had improved insulin sensitivity relative to Control fed mice after 22 months on diet (**Fig. 3T**) but not at other time points (**Figs. S3A-B**), whereas females showed no improvements in insulin sensitivity, despite improvements in glucose tolerance (**Fig. 3U**).

### Isoleucine restriction increases food intake and energy expenditure in old HET3 mice

As with young mice, old male mice on a Low Ile and Low AA diet consumed significantly more calories than Control fed mice (**Fig. 4A**). Despite this increase, isoleucine consumption remained significantly lower in Low Ile-fed mice than Control fed mice (**Fig. 4B**). Across the lifespan study, Low Ile and Low AA-fed mice always ate at least as many calories as Control-fed mice and were at no time calorie restricted (**Figs. 4C**). Old female mice showed a similar pattern, with Low AA and Low Ile mice showing greater calorie consumption, while Low Ile-fed mice maintained significantly reduced isoleucine consumption (**Figs. 4D-E**). As with males, lifespan study, Low Ile and Low AA-fed mice always ate at least as many calories as Control-fed mice and were at no time calorie restricted (**Fig. 4F**).

**Figure 4:**
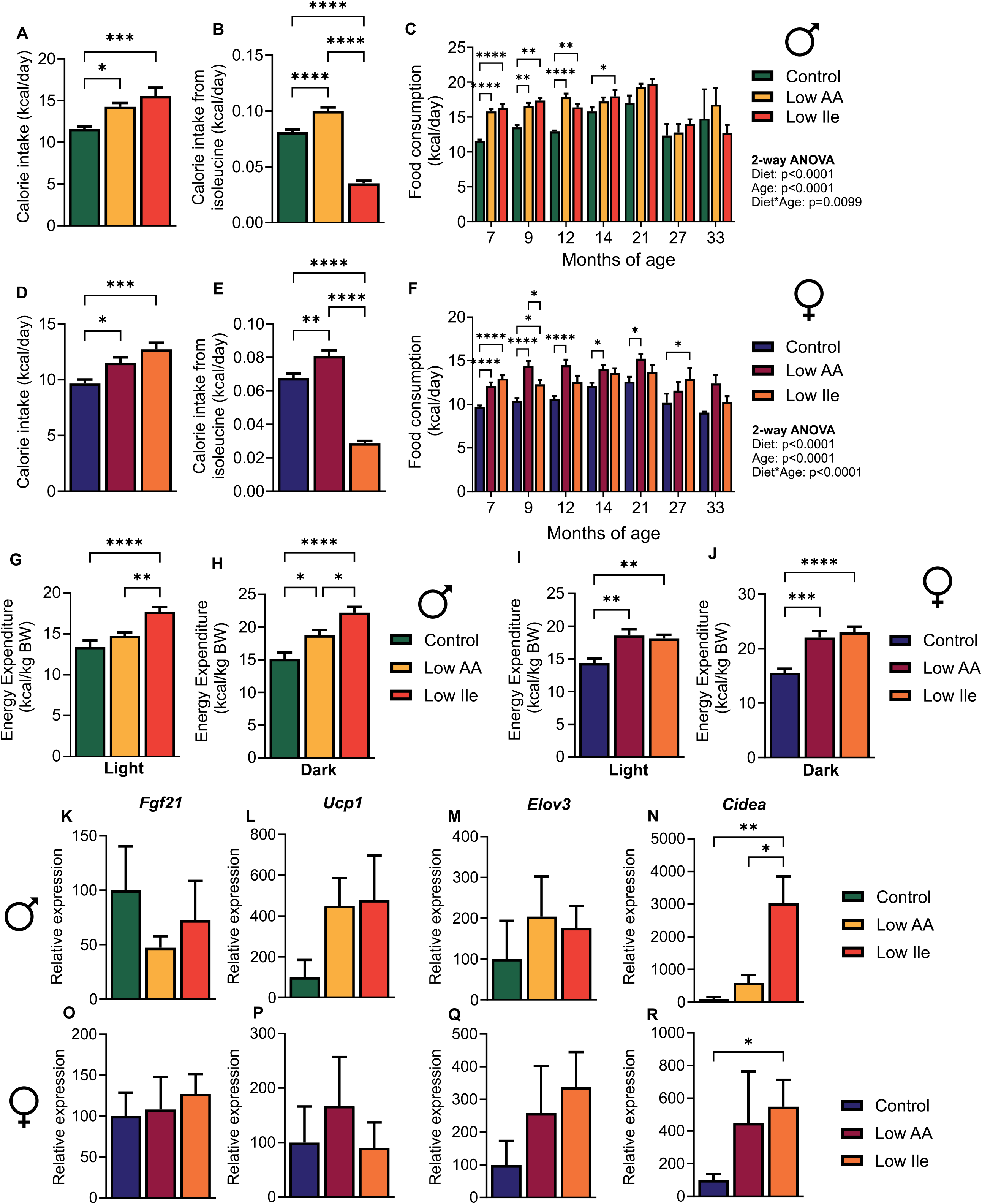
A Low Ile diet started in mid-life increases food consumption and energy expenditure in male and female HET3 mice. (A) Food consumption was measured in home cages after 1 month on diet in male mice. (B) Calculated isoleucine intake for male mice. (C) Food consumption for male mice over the course of the experiment. (D) Food consumption was measured in home cages after 1 month on diet in female mice. (E) Calculated isoleucine intake for female mice. (F) Food consumption for female mice over the course of the experiment. (G-H) Energy expenditure measured in the light and dark phases in male mice. (I-J) Energy expenditure measured in the light and dark phases in female mice. (K-P) Expression of the indicated genes in the iWAT of 24-month-old male (K-M) and female (N-P) mice. (A,B,D,E,G-P) One-way ANOVA for diet with post-hoc Tukey test for pairwise comparisons, (C, F) Two-way ANOVA for Age and Diet with post-hoc Tukey test for multiple comparisons, \**P*<0.05, \*\**P*<0.01, \*\*\**P*<0.001 and \*\*\*\**P*<0.0001. P-values for the overall effect of Diet, Age and the interaction represent the significant p-values from the two-way ANOVA. Data are represented as mean ± SEM.

Due to the prominent weight loss in both males and females on Low Ile diets relative to Control fed mice despite increased food intake, we investigated changes in energy expenditure in these mice. At 24 months of age, we found that Low Ile fed male mice had significantly increased energy expenditure relative to both Control and Low AA fed mice in both the light and dark phase; as expected, we also observed increased energy expenditure in Low AA-fed mice during the dark phase (**Figs. 4G-H**). Both Low AA and Low Ile-fed female mice had increased energy expenditure relative to Control-fed mice in both the light and dark phase (**Figs. 4I-J**).

PR promotes beiging of white adipose tissue in C57BL/6J mice, and we previously observed a similar effect in Low Ile-fed C57BL/6J mice (Hill et al., 2017; Laeger *et al*., 2014; Yu *et al*., 2021). Consistent with white adipose tissue beiging, Low Ile and Low AA-fed male mice showed increased expression of *Ucp1*, although this effect did not reach statistical significance, and increased expression of *Cidea*, but not of *Elov3* or *Fgf21* (**Figs. 4K-N**). In females, *Ucp1* and *Fgf21* were not affected by diet, and although *Elov3* levels were increased, it did not reach statistical significance; however, *Cidea* was significantly upregulated in Low-Ile fed female mice (**Figs. 4O-R**).

### Changes in metabolic health relative to isoleucine intake vary between sexes and with age

We used multivariate analysis to comprehensively identify age and sex-dependent responses to reduced dietary isoleucine. We determined the isoleucine intake of each individual mouse and correlated it with 26 phenotypic measurements obtained from each animal. We used these correlations in a heatmap and then ordered the correlations using hierarchical clustering (**Fig. 5A**). We found that for the most part, old and young males and females had similar phenotypic responses to reduced isoleucine intake. In general, isoleucine intake correlates negatively with energy expenditure, activity and RER and positively with lean mass, body weight, fat mass and glucose area under the curve (poor glycemic control). The strength of the changes varies with age, with old male mice have less strong correlations of isoleucine intake with the measured parameters than young males, and vice versa for females.

**Figure 5:**
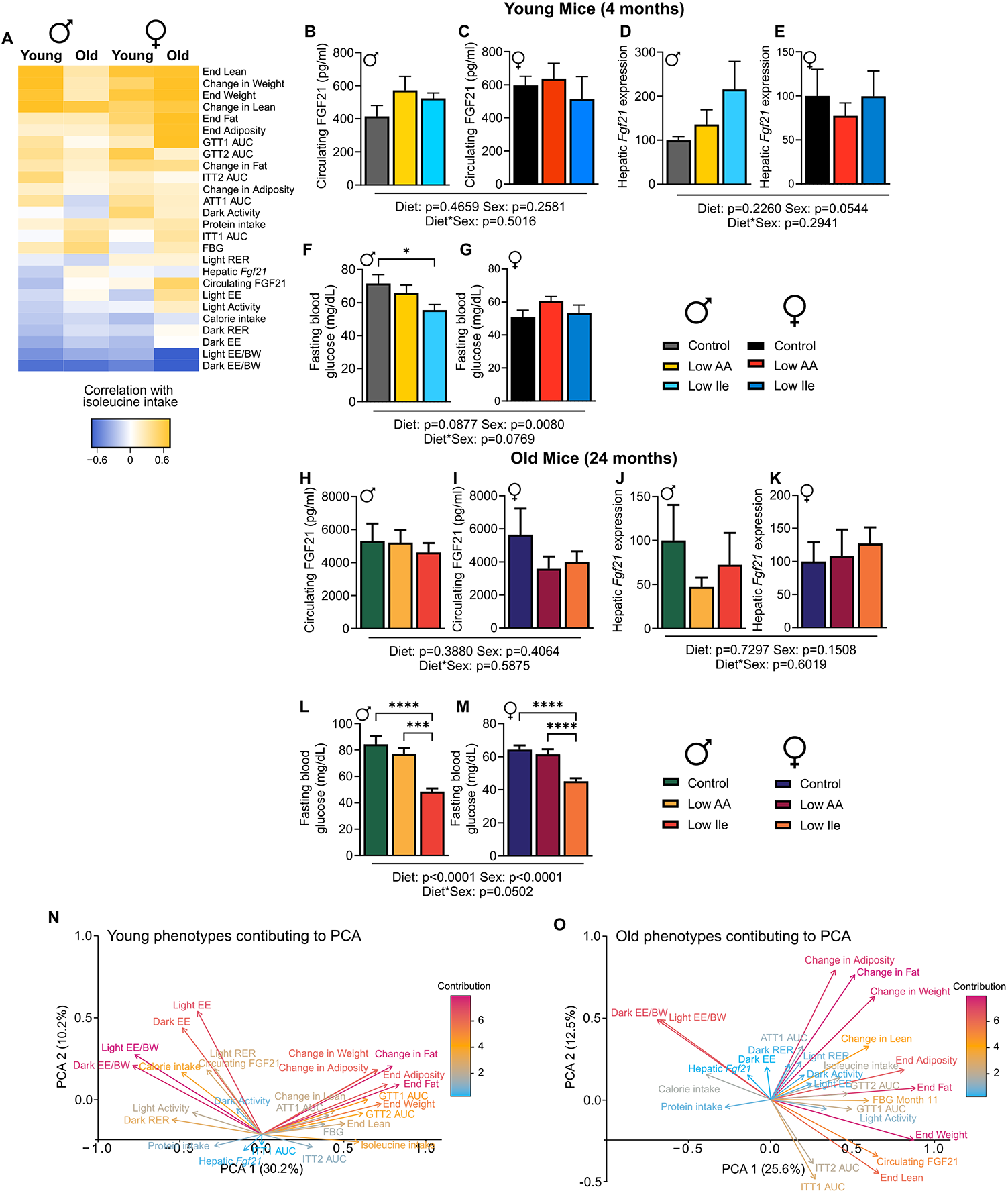
Correlation analysis identifies diet and age dependent and independent physiological and metabolic responses to a Low Ile diet. (A) Phenotypic measurements correlated with consumption of isoleucine (kcals) in each mouse (Pearson’s correlation) and clustered (hierarchical clustering). Phenotypic measurements that do not cluster as well appear in the middle of the correlation plot. (B-M) Selected phenotypic measurements were plotted for young and old, male and female mice. (N-O) Phenotypic measurements from principal component analysis (PCA) of young and old mice were visualized; positively correlated variables point to the same side of the plot, negatively correlated variables point to opposite sides of the plot. Length and color of arrows indicate contribution to the principal components. (B-M) two-way ANOVA between Diet, Sex and the interaction post-hoc Tukey testing for pairwise comparisons; \**P*<0.05, \*\**P*<0.01, \*\*\**P*<0.001 and \*\*\*\**P*<0.0001. P-values for the overall effect of Sex, Diet and the interactions represent the significant p-values from the two-way ANOVA. Data are represented as mean ± SEM.

Interestingly, some measurements such as circulating FGF21 (**Figs. 5B-C & H-I**), hepatic *Fgf21* expression (**Figs. 5D-E & J-K**) and fasting blood glucose (**Figs. 5F-G & L-M**) did not have a particular pattern across groups. We were surprised to see that in both young and old mice, neither a Low AA or Low Ile diet induced expression of hepatic *Fgf21* or increased blood levels of FGF21. Using principal components analysis (PCA) we found that across ages and sexes the phenotypes we measured explained around ∼40% variation in PC1 and PC2 (**Figs. S4A-D**). In young mice, the Low AA group seemed to overlap with Control and Low Ile fed mice; however, in older mice it appeared that Low Ile and Low AA diets shared more similarities (**Figs. S4A-D**). In young mice, differences between groups seem to be particularly driven by changes in energy expenditure and changes in adiposity contributed the most (and oppositely) to the differences in phenotypic variation (**Fig. 5N**). These contributions were also seen in the old mice PCA plot although the relationships were different, with circulating FGF21 contributing to the same direction as energy expenditure in young mice but correlating with insulin sensitivity and weight in old mice (**Fig. 5O**).

### Isoleucine restriction increases lifespan and healthspan in HET3 mice

As mice and humans age they become increasing frail, and we utilized a recently developed mouse frailty index to examine how diet, age, and sex impact frailty (Whitehead et al., 2014). As expected, we observed increased frailty with age in Control-fed mice of both sexes (**Figs. 6A-B**). Overall, frailty scores were significantly lower in older Low AA and Low Ile-fed male mice (**Fig. 6A**), as well as in female mice at 22 and 24 months of age (**Fig. 6B**). In addition to frailty, lower urinary track dysfunction increases, particularly in male mice, with age. Using void spot assays as a measure of urinary frequency, we found that urinary spotting was significantly increased in aged male mice on a Control diet relative to mice on Low AA and Low Ile diets; we observed no differences with diet in aged female mice (**Figs. 6C-D**).

**Figure 6:**
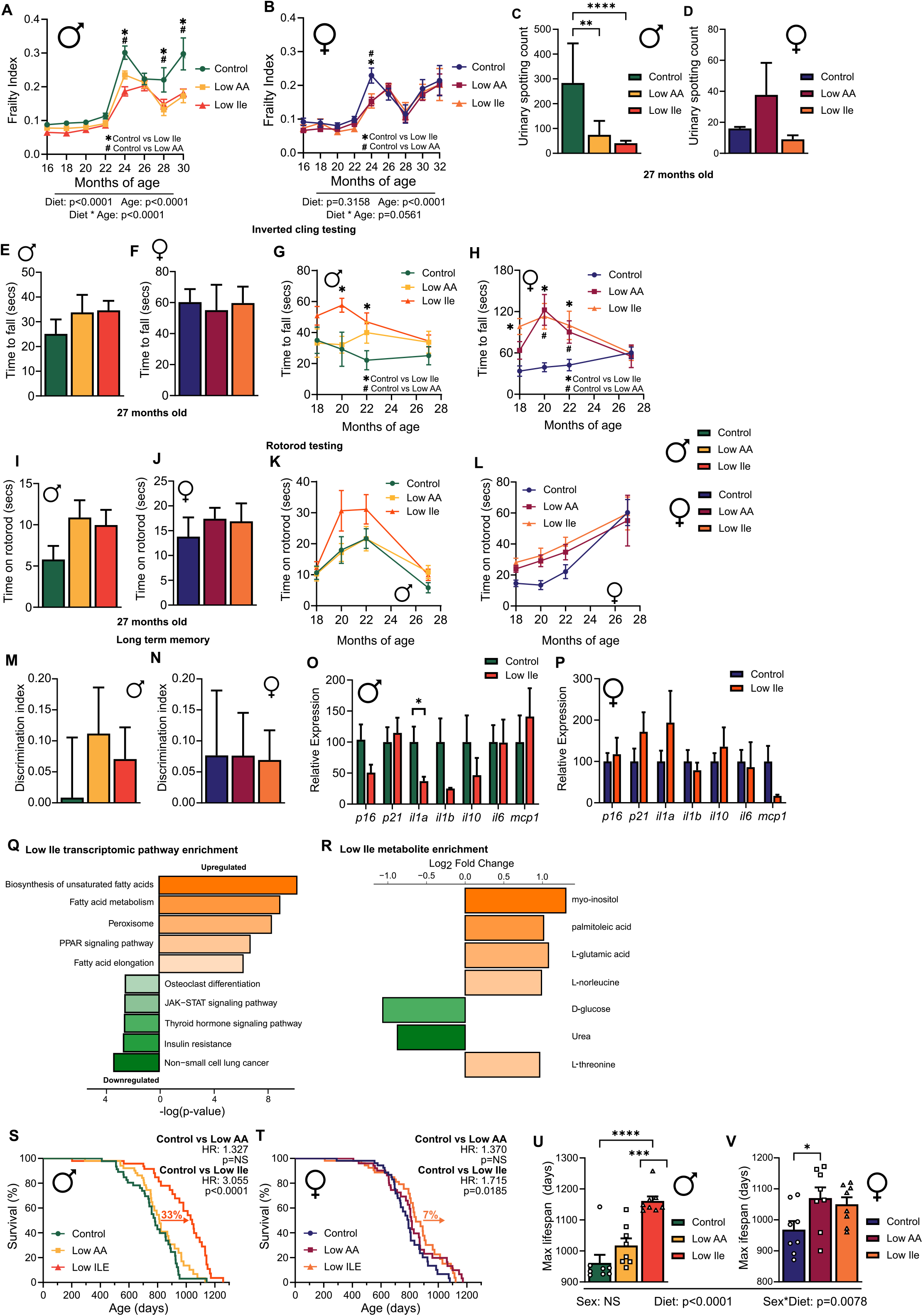
A Low Ile diet increases lifespan and improves healthspan in HET3 male mice. (A-B) Frailty assessments for male (A) and female (B) mice over time. (C-D) Void spot assay was conducted in 27-month-old male (C) and female (D) mice. (E-F) Inverted cling test in 27-month-old male (E) and female (F) mice. (G-H) Inverted cling tests over time in male (G) and female (H) mice. (I-J) Rotorod test in 27-month-old male (I) and female (J) mice. (K-L) Rotorod tests over time in male (K) and female (L) mice. (M-N) Novel object recognition of long-term memory in male (M) and female (N) mice. (O-P) Hepatic expression levels of the indicated genes in 24-month-old male (O) and female (P) mice. (Q) Top significantly enriched transcriptomic pathways in male 24-month-old Control vs Low Ile mice using KEGG. (R) Log2 fold-changes for significantly differentiated metabolites between Control vs Low Ile 24-month-old male mice. (S) Survival curves by diet in male mice. (T) Survival curves by diet in female mice. (U) Average top 10% of lifespans in male mice by diet. (V) Average top 10% of lifespans in female mice by diet. (A-B) Mixed-effects model (REML) for time and diet with post-hoc Tukey testing for multiple comparisons. (C-F, I-J, M-N, Q-R) One-way ANOVA for diet with post-hoc Tukey test for pairwise comparisons; \**P*<0.05, \*\**P*<0.01, \*\*\**P*<0.001 and \*\*\*\**P*<0.0001. (O-P) Log-rank (Mantel-Cox) test for Control vs Low AA and Control vs Low Ile. Data are represented as mean ± SEM.

We also investigated strength in mice over time using inverted cling assay and rotorod testing, as dietary protein and BCAAs are commonly associated with muscle strength, and dietary protein is often recommended to the elderly who are at risk from sarcopenia. In late life, we found no significant differences in inverted cling performance between the diet groups in either males or females (**Figs. 6E-F**). In younger mice of both sexes, mice consuming a Low Ile diet were able to cling for longer than Control-fed mice; a similar affect was observed in Low AA-fed female mice (**Figs. s6G-H**). In both sexes, this effect disappeared with age, concurrently with the convergence of body weights (**Fig. 6H**). Similarly, with rotorod testing we did not find significant differences between the diet groups in aged males or females (**Figs. 6I-J**) and we saw no significant differences at any time points (**Figs. 6K-L**). We also investigated changes in memory on different diets and found no changes in either short-term or long-term memory through novel object recognition tests (**Figs. 6M-N**).

To investigate if senescence was reduced in Low Ile males we measured the expression of several senescence-associated secretory phenotype (SASP) markers in the liver of Low Ile fed mice relative to Controls. We found that, in males only, the inflammatory marker interleukin 1-α (*Ila*) was downregulated, and no changes were seen in the markers measured in females (**Figs. 6O-P**). We also examined transcriptomic changes in the liver and found that in Low Ile fed males relative to Control males, several pathways related to changing fatty acid metabolism were significantly upregulated, while immune and insulin resistance pathways were significantly downregulated (**Fig. 6Q**). In addition, when exploring changes in hepatic metabolism, we found that Low Ile fed males showed increases in the omega-7 monounsaturated fatty acid palmitoleic acid and the amino acids L-glutamic acid and L-threonine but reduced levels of D-glucose and urea (**Fig. 6R**).

In agreement with the significance difference in frailty, we found that consumption of a Low Ile diet from 6 months of age significantly increased lifespan, with a 33% increase in median lifespan relative to Control-fed mice (**Fig. 6S**). Somewhat surprisingly, despite inhibiting frailty, a Low AA diet had no effect on median lifespan (**Fig. 6S**). A Low Ile diet also increased median lifespan in females relative to Control-fed mice by a more modest 7%; once again, there was no effect of a Low AA diet on median lifespan (**Fig. 6T**). We also observed a significant effect of diet on maximum lifespan (90%); in males a Low Ile diet increased maximum lifespan relative to both Low AA-fed (by 13%) and Control-fed (by 17%) fed mice (**Fig. 6U**). In females, a Low AA diet significantly increase maximum lifespan relative to Control-fed mice by 12% (**Fig. 6V**).

## Discussion

Recently it has become clear that the macronutrient composition of the diet is a key factor in the “food as medicine” approach to health (Green and Lamming, 2021). Multiple studies have highlighted dietary protein as a key determinant of human health (Ferraz-Bannitz *et al*., 2022; Fontana *et al*., 2016; Lagiou et al., 2007; Levine et al., 2014; Sluijs et al., 2010; Vergnaud et al., 2013). In animals as well, dietary protein has identified as a key regulator of metabolic health as well as longevity (Solon-Biet *et al*., 2014; Solon-Biet et al., 2015).

We and others have hypothesized that many of the beneficial effects of PR may result from the reduced consumption of specific AAs. We have previously shown that 67% restriction of the three BCAAs is sufficient to recapitulate the benefits of a LP diet, and can extend the lifespan of male mice equivalently to an equivalent restriction of all twenty common amino acids (Richardson *et al*., 2021). The three BCAAs do not all have equivalent metabolic effects, and we have shown that isoleucine in particular is a key regulator of metabolic health. Restriction of isoleucine is both necessary for the metabolic benefits of a LP diet, and is sufficient to promote glucose tolerance, hepatic insulin sensitivity, energy expenditure and to reverse the effects of Western diet-induced obesity on body composition, glycemic control, and hepatic steatosis (Yu *et al*., 2021).

Here, we tested the hypothesis that restriction of isoleucine alone would promote health span and longevity. As our recent work has shown that sex and genetic background on the beneficial effects of LP diets (Green *et al*., 2022), we chose to investigate this in genetically heterogenous HET3 mice. We find that, even in a heterogeneous population, a Low Ile diet has dramatic effects on weight, fat mass, glycemic control and energy expenditure in both young and old mice of both sexes. Interestingly, these effects were distinct – and generally greater in magnitude – than the effects in mice consuming a Low AA diet in which all amino acids were restricted. Furthermore, we found that in males, a low isoleucine diet is able to substantially increase both median and maximum lifespan while reducing frailty; a Low Ile diet had a more modest effect on the median lifespan of female HET3 mice, while females fed a Low AA diet did not have increased median lifespan but did have a significantly increased maximum lifespan.

When started in young mice, a Low Ile diet significantly improves metabolic health in both HET3 male and female life. Our findings here mirror our previous work, where found a similar pattern of weight loss in C57BL/6J males fed a Low Ile diet (Yu *et al*., 2021). HET3 mice on a Low AA diet also showed a modest reduction in body weight relative to Control fed mice, similar to our previous work with HET3 mice on protein restriction (Green *et al*., 2022). Importantly, when started in adulty mice, a Low Ile and to a lesser extent a Low AA diet was still able to promote weight loss in male and female HET3 mice. Interestingly, the Low Ile diet had a much greater impact on glucose tolerance than the Low AA diet under most conditions examined. All of this highlights the potential translatability of an isoleucine-restricted diet.

A potential caveat of nutritional intervention, particularly those initiated later in life, is the potential impact of a restricted diet on sarcopenia (Ham et al., 2022). Comfortingly, in humans there is no relationship between intake of total or individual BCAAs and sarcopenia (Ebrahimi-Mousavi et al., 2022). Our work here found that a Low Ile diet tended to promote leanness, and grip strength (inverted cling test) and rotorod testing did not show any significant defect in either Low AA or Low Ile groups. While grip strength appeared to be improved in middle age for Low Ile-fed mice, this is most likely due to reduced body weight. However, we did not examine muscle strength and quality in great detail, and more directly assessing the effects of dietary isoleucine on muscle strength, quality and fiber type would be useful for a future study.

Many of the metabolic benefits of a LP diet have been linked to LP-induced expression of the energy balance hormone FGF21; in absence of FGF21, many of the beneficial effects of a LP diet are ablated (Green *et al*., 2022; Hill et al., 2022; Hill *et al*., 2017; Hill et al., 2019; Laeger *et al*., 2014). However, despite observing effects on body weight and composition, food consumption, energy expenditure, glucose tolerance and changes in fatty acid metabolism – effects which have been linked in part to FGF21 in previous studies - we saw no changes in either circulating FGF21 or hepatic *Fgf21* expression in mice fed either Low AA or Low Ile diets. These results are in accordance with a study of natural sourced low protein diets we recently conducted, in which we observed that FGF21 was induced by PR in C57BL/6J mice, but not in HET3 or DBA/2J mice (Green *et al*., 2022). As previous studies, which have convincingly identified a role for FGF21 in the response to PR, have been conducted principally in male C57BL/6J mice, perhaps induction of FGF21 is not required for the effects of PR in other genetic background. Another possibility is that Low AA or Low Ile diets may have induced FGF21 for a short period of time, after which time the levels in these mice returned to the same level found in the Control mice; or that the levels of FGF21 in the blood much be at a certain level in order to permit Low AA or Low Ile-induced changes, but do not necessarily need to increase. Understanding the role of FGF21 should be a priority for future studies.

Notably, restriction of many individual amino acids, including methionine, leucine, isoleucine, histidine, threonine, tryptophan, and valine as well as simultaneous restriction of the three BCAAs, have been associated with increased energy expenditure, at least in C57BL/6J males, but not all of these restrictions induce the FGF21-UCP1 thermogenic axis in iWAT (Cummings et al., 2018; Flores et al., 2022; Fontana *et al*., 2016; Forney et al., 2020; Lees et al., 2017; Yap et al., 2020; Yu *et al*., 2021). Similarly, in HET3 mice fed either the Low AA or Low Ile diets, there is not a clear upregulation of iWAT *Ucp1* in males and no change in females. Interestingly, we did see a statistically significant increase in *Cidea*, but not *Elov3* or *Ucp1* in the iWAT of both male and female Low Ile-fed mice. *Cidea* has a complex set of interactions; while *Cidea* deficient mice are lean and resistant to obesity, it has also been suggested that *Cidea* may regulate lipolysis and thermogenesis through *Ucp1* suppression (Zhou et al., 2003). In humans, reduced *CIDEA* expression is associated with high body fat and a low basal metabolic rate, and is increased in adipose tissue by a calorie restriction (Gummesson et al., 2007). Future work will need to be conducted to determine if induction of *Cidea* contributes to the increased energy expenditure and other metabolic benefits of a Low Ile diet.

Our most striking finding is the robust extension of both median and maximum lifespan induced in HET3 male mice by feeding of a Low Ile diet beginning at 6 months of age. The ability of isoleucine restriction to extend the lifespan of this heterogeneous population, as well as the effect of isoleucine restriction on frailty and maximum lifespan, clearly indicate that the benefits of isoleucine restriction are mediated by geroprotective effects. due to impacts of strain-specific causes of death. The effect of isoleucine restriction on median female lifespan was more modest, and the effect on maximum lifespan did not reach statistical significance. This male-specific or male-biased effect has been seen before in a number of interventions by the National Institute on Aging Interventions Testing Program (Harrison et al., 2014; Strong *et al*., 2016). However, it is notable that pharmacological or genetic disruption of the amino acid sensitive mTORC1 signaling pathway has larger benefits for the lifespan of female mice than male mice (Lamming, 2014; Lamming et al., 2012; Miller et al., 2014; Selman et al., 2008). This suggests that isoleucine restriction may either work through largely mTORC1-independent mechanisms, or that – as we found with BCAA restriction (Richardson *et al*., 2021) – isoleucine restriction inhibits mTORC1 signaling preferentially in males.

Intriguingly, we did not observe a significant increase in either median or maximum lifespan in Low AA-fed male HET3 mice, although we did observe an increased in maximum lifespan in Low AA-fed female mice. This is despite the fact that we and other groups have reported that an LP diet increases the lifespan in C57BL/6J male mice, and we have shown that a Low AA diet increased the lifespan of C57BL/6J mice by over 30% (Hill *et al*., 2022; Richardson *et al*., 2021; Solon-Biet *et al*., 2014). In addition to the genetic background, another difference between these previous studies and the results reported here is that we initiated the diets later than these previous studies – and studies in which restriction of protein or amino acids were done earlier tend to have reported larger effects than studies initiating restriction later in life. Similar effects have been reported for CR (Hahn et al., 2019), and it is possible that we initiated LP diet feeding too late to see robust effects on lifespan.

In terms of improving healthspan, Low Ile and Low AA diets had by far the greatest effect on healthspan (through reduced frailty and bladder dysfunction) in male HET3 mice, with a much smaller effect on female frailty. This agrees generally with our finding that a Low BCAA diet improves frailty only in male, not female, C57BL/6J mice (Richardson *et al*., 2021). Importantly, it also demonstrates that the clinical frailty index for mice is applicable not only to inbred strains, but also to genetically heterogeneous mice.

In conclusion, we have shown that dietary restriction of a single branched-chain amino acid, isoleucine, can extend both healthspan and lifespan when begun at 6 months of age – roughly equivalent to a human in their 30’s (Flurkey *et al*., 2010). This effect is particularly robust in males, but benefits are observed in both sexes; and the fact that isoleucine restriction works in a genetically heterogeneous population suggests that such an intervention may be applicable to humans as well. Additional research will be required to determine if there are potentially negative effects of isoleucine restriction, and to examine how optimal levels of isoleucine for lifespan and health span vary with age and sex. Our results demonstrate that in agreement with emerging human data (Deelen *et al*., 2019; Yu *et al*., 2021), isoleucine is critically important in metabolic health and aging, and provide additional evidence that protein quality – the specific AA composition of dietary protein – is as important, or even more important, than the amount of protein consumed. While additional research and randomized clinical trials will be required to determine how dietary isoleucine affects healthy aging in humans, and to determine if limiting isoleucine is practical in the clinical setting, our results support an emerging consensus that limiting dietary levels of specific essential amino acids may provide the secret to a long and healthy life.

## Limitations of study

Limitations of our work include that we examined only a single level of dietary restriction. Further, the diets used here are based on the amino acid profile of whey rather than casein; however, LP studies conducted with either protein source have found broadly similar effects (Fontana *et al*., 2016; Laeger *et al*., 2014; Maida et al., 2017; Maida et al., 2016; Solon-Biet *et al*., 2014; Yu *et al*., 2021). To keep diets isocaloric, the reduction of dietary amino acids from the Low AA diet was balanced by addition of carbohydrates, and the reduction of isoleucine from the Low Ile diet was balanced with non-essential amino acids. The protein to carbohydrate ratio, the type of dietary carbohydrate consumed, the precise degree of restriction, the specific non-essential amino acids added to the Low Ile diet, and diet-induced changes to the microbiome could all play a role in the responses we observe here (MacArthur et al., 2022; Pak et al., 2019; Solon-Biet *et al*., 2014; Solon-Biet *et al*., 2015; Wali et al., 2021). Finally, our molecular analysis was largely limited to the liver, and while this is the first organ to be exposed to absorbed nutrients, dietary isoleucine metabolism occurs in many tissues (Neinast et al., 2019).

## Materials and Methods

### Mouse information

All procedures were performed in conformance with institutional guidelines and were approved by the Institutional Animal Care and Use Committee of the William S. Middleton Memorial Veterans Hospital (Madison, WI, USA).

HET3 mice are the F2 progeny of (BALB/cJ x C57BL/6J) mothers and (C3H/HeJ x DBA/2J) fathers; female BALB/cJ (#000651), male C57BL/6J (#000664), female C3H/HeJ (#000659) and male DBA/2J (#000671) were obtained from The Jackson Laboratory and bred to produce heterogeneous HET3 F2 mice. Mice were acclimatized on chow diet Purina 5001 for one week before experiment start and housed 2-3 per cage. All mice were maintained at a temperature of approximately 22°C, and health checks were completed on all mice daily. The short-term study was started when the mice were 9 weeks of age, and the lifespan study when they were 6 months old.

At the start of the experiment, mice were randomized to receive either the 22% (Control, TD.140711) or 7% (Low AA, TD.140712) amino acid diet, or 22% amino acid diet with isoleucine reduced by 2/3rds (Low Ile, TD.160734); all diets were obtained from Envigo. Within the diet series, calories from amino acids were replaced by calories from carbohydrates, while calories from fat were held fixed at 20%, making the diets isocaloric (3.6 Kcal/g). For the Low Ile diet, amino acid levels were made up with non-essential amino acids, making the diet isonitrogenous with the Control diet. Full diet descriptions are provided in **Table S1**. The randomization of mice was performed at the cage level to ensure that all groups had approximately the same initial starting weight and body composition. The number of animals in each group used for each experiment is listed in **Table S2** and the N for each figure is listed in **Table S3**. Mice were housed in a SPF mouse facility in static microisolator cages, except when temporarily housed in a Columbus Instruments Oxymax/CLAMS metabolic chamber system. Mice were housed under a 12:12 h light/dark cycle with free access to food and water, except where noted in the procedures below.

### In vivo Procedures

Glucose, insulin, and alanine tolerance tests were performed by fasting the mice overnight for 16 hours and then injecting glucose (1 g kg^−1^), insulin (0.75 U kg^-1^) or alanine (2 g kg^-1^) intraperitoneally (i.p.) (Bellantuono et al., 2020; Yu et al., 2018). Glucose measurements were taken using a Bayer Contour blood glucose meter (Bayer, Leverkusen, Germany) and test strips.

Mouse body composition was determined using an EchoMRI Body Composition Analyzer (EchoMRI, Houston, TX, USA). For assay of multiple metabolic parameters [O2, CO2, food consumption, respiratory exchange ratio (RER), energy expenditure] and activity tracking, mice were acclimated to housing in a Oxymax/CLAMS metabolic chamber system (Columbus Instruments) for ∼24 h and data from a continuous 24 h period was then recorded and analyzed. Food consumption in home cages was measured by moving mice to clean cages, filling the hopper with a measured quantity of fresh diet in the morning and measuring the remainder in the morning 3-6 days later. The amount was adjusted for the number of mice per cage, the number of days that passed and the relative weights of the mice (i.e., heavier mice were credited with a larger relative portion of the food intake). Mice were euthanized by cervical dislocation after an overnight (16h) fast and tissues for molecular analysis were flash-frozen in liquid nitrogen or fixed and prepared as described below.

### Void Spot Assay

Void spot assays were performed as described previously (Keil et al., 2016). Briefly, mice were individually placed in standard mouse cages with thick chromatography paper (Ahlstrom, Kaukauna, WI). During the 4 hour study period, mice were restricted from water. Chromatography papers were imaged with a BioRad ChemiDoc Imaging System (BioRad, Hercules, CA) using an ethidium bromide filter set and 0.5 second exposure of ultraviolet light. Images were imported into ImageJ and total void spots analyzed with VoidWhizzard (Wegner et al., 2018).

### Assays and Kits

Blood for circulating FGF21 analysis was obtained following an overnight fast. Blood FGF21 levels were assayed by a mouse/rat FGF-21 quantikine ELISA kit (MF2100) from R&D Systems (Minneapolis, MN, USA).

### Quantitative PCR

Liver was extracted with Trireagent (Sigma, St Louis, MO, USA). Then, 1 μg of RNA was used to generate cDNA (Superscript III; Invitrogen, Carlsbad, CA, USA). Oligo dT primers and primers for real-time PCR were obtained from Integrated DNA Technologies (IDT, Coralville, IA, USA). Reactions were run on a StepOne Plus machine (Applied Biosystems, Foster City, CA, USA) with Sybr Green PCR Master Mix (Thermo Fisher Scientific, Waltham, MA). Actin was used to normalize the results from gene-specific reactions. Primer sequences can be found in **Table S4**.

### Transcriptomic Analysis

RNA was extracted from liver as previously described (Cummings *et al*., 2018). The concentration and purity of RNA was determined using a NanoDrop 2000c spectrophotometer (Thermo Fisher Scientific, Waltham, MA) and RNA was diluted to 100-400 ng/µl for sequencing. The RNA was then submitted to the University of Wisconsin-Madison Biotechnology Center Gene Expression Center & DNA Sequencing Facility, and RNA quality was assayed using an Agilent RNA NanoChip. RNA libraries were prepared using the TruSeq Stranded Total RNA Sample Preparation protocol (Illumina, San Diego, CA) with 250ng of mRNA, and cleanup was done using RNA Clean beads (lot #17225200). Reads were aligned to the mouse (*Mus musculus*) with genome-build GRCm38.p5 of accession NCBI:GCA_000001635.7 and expected counts were generated with ensembl gene IDs (Zerbino et al., 2018).

Analysis of significantly differentially expressed genes (DEGs) was completed in R version 3.4.3 (Team, 2017) using *edgeR* (Robinson et al., 2009) and *limma* (Ritchie et al., 2015). Gene names were converted to gene symbol and Entrez ID formats using the *mygene* package. To reduce the impact of external factors not of biological interest that may affect expression, data was normalized to ensure the expression distributions of each sample are within a similar range. We normalized using the trimmed mean of M-values (TMM), which scales to library size. Heteroscedasticity was accounted for using the voom function, DEGs were identified using an empirical Bayes moderated linear model, and log coefficients and Benjamini-Hochberg (BH) adjusted p-values were generated for each comparison of interest (Benjamini and Hochberg, 1995). DEGs were used to identify enriched pathways, both Gene Ontology (for Biological Processes) and KEGG enriched pathways were determined for each contrast. All genes, log_2_ fold-changes and corresponding unadjusted and Benjamini-Hochberg adjusted p-values can be found in **Table S5**.

### Metabolomic Analysis

Untargeted metabolomics was performed as previously described (Fiehn, 2016). Samples were kept at -20°C throughout the extractions in an Iso-Therm System (VWR #20901-646). Fifteen milligrams of liver were homogenized in 200 µL of a 3:3:2 solution of acetonitrile:isopropanol:water (MeCN:IPA:H_2_O) in ceramic bead tubes (1.4 mm, Qiagen #13113-50) using a TissueLyzer II (Qiagen #85300). An additional 800 µL of the extraction solvent was added and samples were centrifuged for 2 min at 14,000xg at RT. Lysate equivalent to 2 mg of tissue was transferred to a new 1.5 mL tube and 100 µL of extraction solvent containing 5 ug succinate-d4 (Sigma-Aldrich #293075) and 1 µg myristate-d27 (CDN Isotopes #D-1711) internal standards was added to each sample to a final volume of 500 uL. 250 uL of extract was transferred to a new tube and dried down in a SpeedVac Plus SC110A (), then resuspended in 420 uL 1:1 MeCN:H_2_O. Samples were centrifuged as before and 400 µL was transferred to a glass autosampler vial with insert and dried down using a SpeedVac. To the dried extract, 10 uL of 20 mg/mL methoxyamine hydrochloride (MP Biomedicals #155405) in pyridine (Millipore PX2012-7) was added and vials were capped, then incubated at 37°C for 1.5h with light flicking every 30 min to mix. 91 µL of MSTFA (Thermo Fisher Scientific #TS-48915) was added to vials and incubation at 37°C with constant shaking for an additional 30 min was done to derivatize samples. Finally, samples were transferred to a GC-MS vial and immediately queued for injection.

Analysis of trimethylsilylated metabolites was performed using an Agilent 5977A Series GC-MS. A Phenomenex ZB-5MSi column (30 m, 0.25 I.D., 0.25 um; #7HG-G018-11) with 5 m column guard was used for chromatographic separation. Splitless, 1 µL sample injections were performed with injector port at 250°C with a 60 s purge at 8.2 psi. Helium was used as carrier gas and kept at 1 mL/min during the GC profile which was as follows: oven at 60°C for 1 min followed by an increase of 10°C/min to 325°C, which was held for 10 min. The MS transfer line was kept at 290°C with a solvent delay of 5.70 min. Ion source was kept at 230°C, quadrupole at 150°C and the acquisition range was from 50-750 m/z collecting 3 spectra/sec. Two quality control samples of pooled liver extracts were run at the beginning and end of each day to monitor for changes in chromatography and to equilibrate the system. A FAME mix was run externally for RI calibration. Collected data was processed in the NIST AMDIS Program using the Fiehn metabolomics library (Kind et al., 2009) for compound identification and peak integration before being exported to R for normalization to the succinate-d4 internal standard to control for extraction variability. Final data was reported as ion count per 2 mg liver. Data was normalized and analyzed using the *metabolomics* package in R. Data was normalized and analysed using the *metabolomics* package in R. Log2 fold-changes and associated p-values for 24 month old male mice on Low AA or Low Ile diets relative to Control fed mice can be found in **Table S6**.

### Statistical Analyses

Most statistical analyses were conducted using Prism, version 8 (GraphPad Software Inc., San Diego, CA, USA) and R (version 4.1.0). Tests involving multiple factors were analyzed by either a two-way analysis of variance (ANOVA) with Sex and Diet as categorical variables, followed by a Tukey–Kramer *post hoc* test for multiple comparisons or by one-way ANOVA with Diet as the categorical variable followed by a Tukey–Kramer *post hoc* test for multiple comparisons. Data distribution was assumed to be normal but was not formally tested. Transcriptomics and metabolomics data were analyzed using R (version 3.3.1). All correlations where depicted were produced using Pearson’s correlations and PCA plots were produced by imputing missing data and scaling the data using the R package “missMDA” using the PCA analysis and plots were generated using the R package “factoextra” (Kassambara and Mundt, 2020; Lê et al., 2008).

## AUTHOR CONTRIBUTIONS

CLG, IMO, JS, and DWL conceived of and designed the experiments. CLG, MET, RB, RJ, HHP, AB, GN, MMS, C-YY, MFC, VF, TTL, and SN performed the experiments. CLG, RJ, KC, TTL, KAM, IMO, JS, and DWL analyzed the data. CLG, TTL, WAR, KAM, IMO, JS, and DWL secured funding and supervised personnel. CLG, TTL, IMO, JS, and DWL wrote the manuscript.

## ACKNOWLEDGEMENTS

We thank all members of the Lamming lab for their feedback, and Dr. Tina Herfel (Envigo) for assistance with diets. The Lamming lab is supported in part by the NIA (AG056771, AG062328, and AG061635), the NIDDK (DK125859), and startup funds from UW-Madison. CLG was supported in part by Dalio Philanthropies and was a Glenn Foundation for Medical Research Postdoctoral Fellow. HHP was supported in part by F31AG066311. MET was supported in part by a Supplement to Promote Diversity in Health-Related Research (R01AG062328-03S1). RB is supported by training grant T32DK007665. TTL was supported by K01AG059899. The Simcox lab is supported in part by the NIDDK (R01DK133479), a pilot grant to JS from the Diabetes Research Center at Washington University, P30DK020579, and a UW BIRCWH Scholars Program award to JS (K12HD101368). JS is an American Federation for Aging Research (AFAR) grant recipient. IO was supported by UWCCC Support Grant P30 CA014520 and Wisconsin Head and Neck Cancer SPORE CEP P50DE026787. WAR was supported by U54DK104310 and R01DK131175. Support was also provided by the UW-Madison OVCRGE with funding from the Wisconsin Alumni Research Foundation. This work supported in part by the U.S. Department of Veterans Affairs (I01-BX004031), and this work was supported using facilities and resources from the William S. Middleton Memorial Veterans Hospital. The content is solely the responsibility of the authors and does not necessarily represent the official views of the NIH. This work does not represent the views of the Department of Veterans Affairs or the United States Government.

---

**Supplementary Figure 1.**
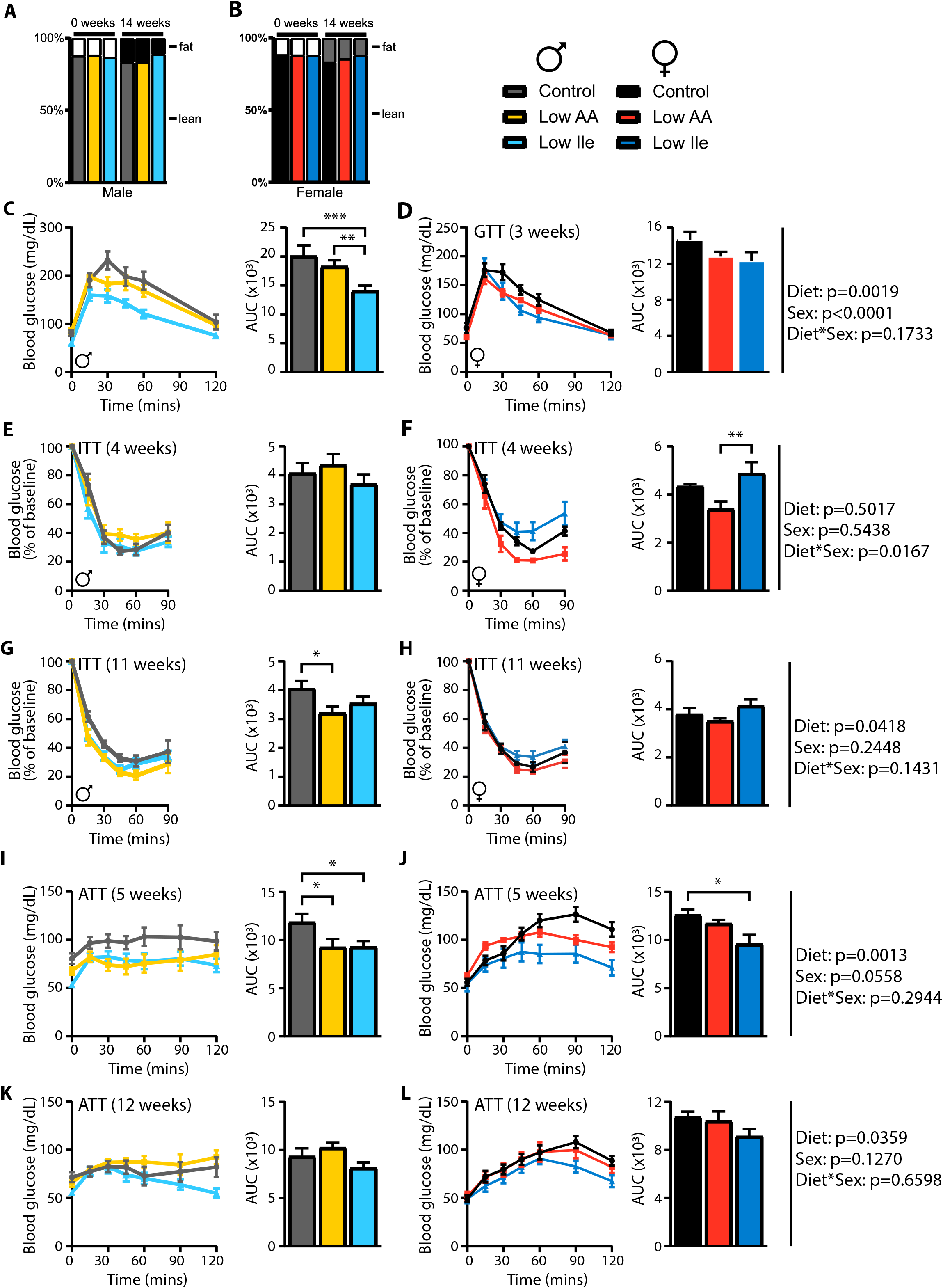
(A) Percentage body composition change from beginning and end of study in males. (B) Percentage body composition change from beginning and end of study in females. (C) Male glucose tolerance test (GTT) after 3 weeks on diet and representative area under the curve (AUC). (D) Female GTT after 3 weeks on diet and representative area under the curve (AUC). (E) Male insulin tolerance test (ITT) after 4 weeks on diet and representative area under the curve (AUC). (F) Female ITT after 4 weeks on diet and representative area under the curve (AUC). (G) Male insulin tolerance test (ITT) after 11 weeks on diet and representative area under the curve (AUC). (H) Female ITT after 11 weeks on diet and representative area under the curve (AUC). (I) Male alanine tolerance test (ATT) after 5 weeks on diet and representative area under the curve (AUC). J) Female ATT after 5 weeks on diet and representative area under the curve (AUC). (K) Male alanine tolerance test (ATT) after 12 weeks on diet and representative area under the curve (AUC). (L) Female ATT after 12 weeks on diet and representative area under the curve (AUC). Two-way ANOVA between Sex and Diet groups with post-hoc Tukey test for pairwise comparisons, \**P*<0.05, \*\**P*<0.01, \*\*\**P*<0.001 and \*\*\*\**P*<0.0001. P-values for the overall effect of Sex, Diet and the interaction represent the significant p-values from the two-way ANOVA. Data are represented as mean ± SEM.

**Supplementary Figure 2.**
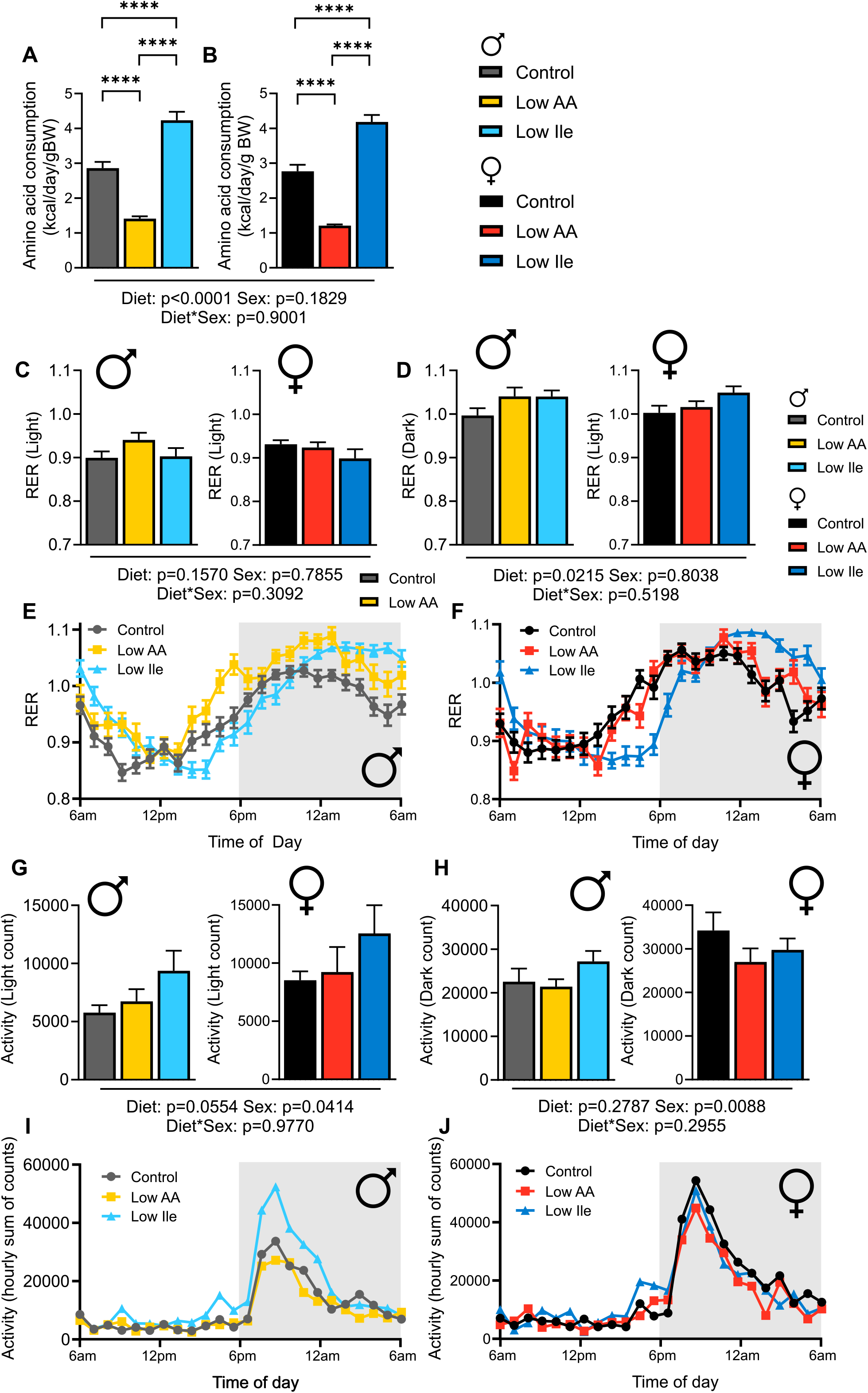
(A) Food consumption was measured in home cages after 3 weeks on diet in male mice and isoleucine intake was calculated for males and (B) females. (C-D) Respectively RER measures in the light and dark phases in male mice and female mice. (E-F) RER of a 24 hour period in male and female mice. (G-H) Activity measured respectively in the light and dark phases in male and female mice. (I-J) Energy expenditure of a 24-hour period in male and female mice. Two-way ANOVA for Diet and Sex with post-hoc Tukey multiple comparison test, \**P*<0.05, \*\**P*<0.01, \*\*\**P*<0.001 and \*\*\*\**P*<0.0001. P-values for the overall effect of Diet, Sex and the interaction represent the significant p-values from the two-way ANOVA. Data are represented as mean ± SEM.

**Supplementary Figure 3.**
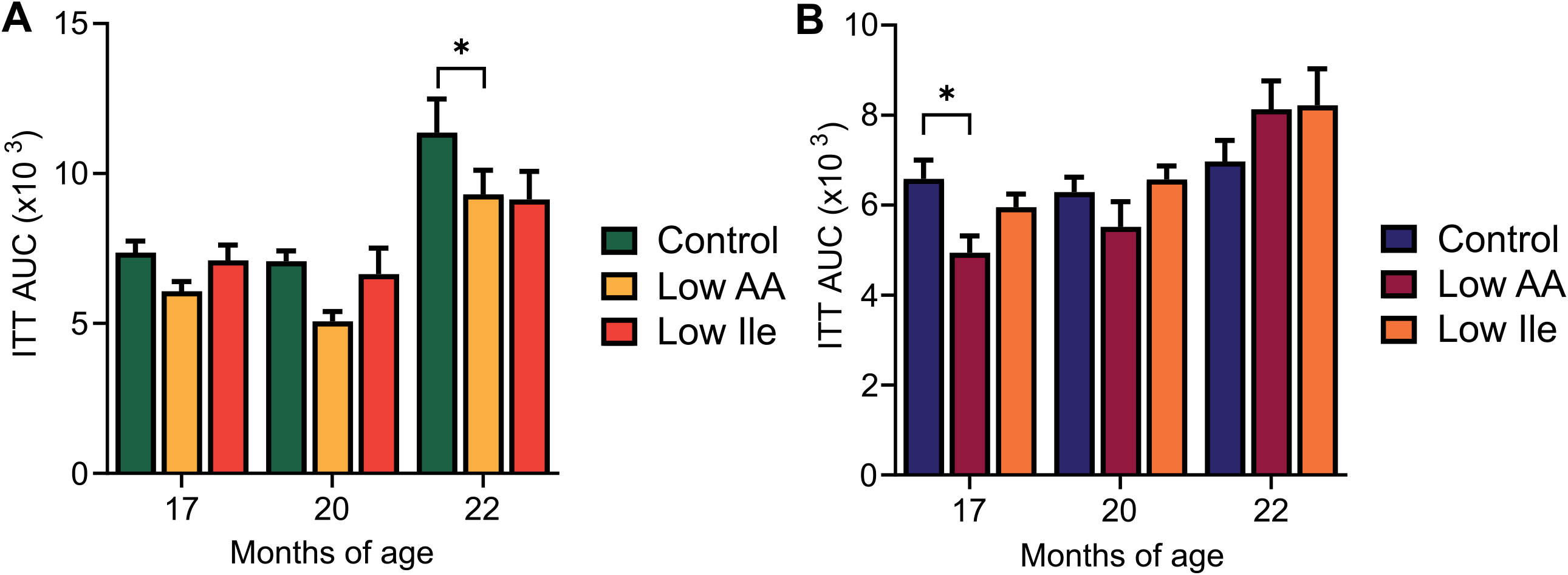
(A) ITT AUC for male mice over the course of the experiment. (B) ITT AUC for female mice over course of experiment. Two-way ANOVA for Diet and Age with post-hoc Tukey multiple comparison test, \**P*<0.05, \*\**P*<0.01, \*\*\**P*<0.001 and \*\*\*\**P*<0.0001.

**Supplementary Figure 4.**
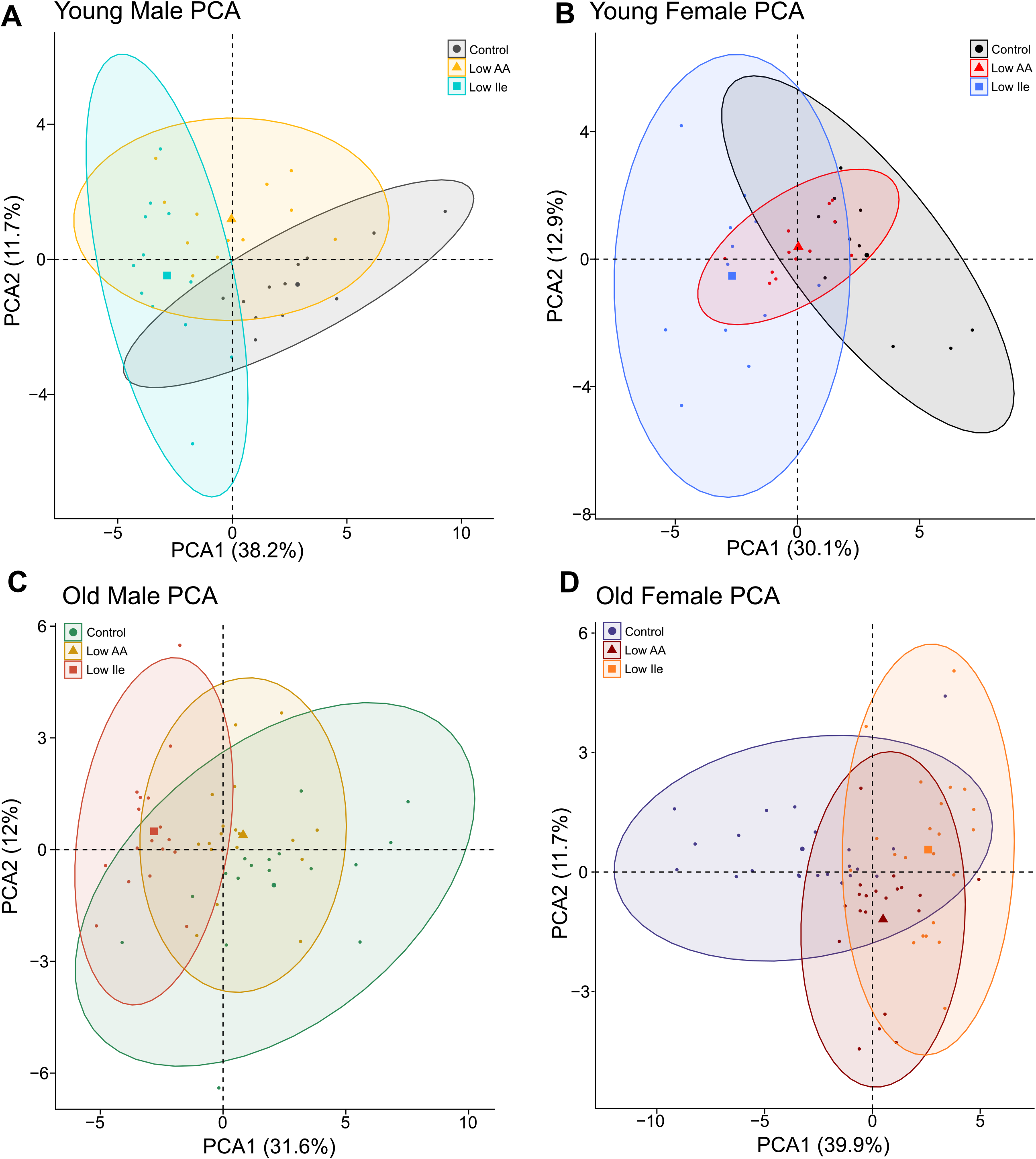
Principal component analysis for 27 measured phenotypes in Young Male Mice (A), Young Female Mice (B), Old Male Mice (C) and Old Female Mice (D).

## Supplemental Tables

**Supplementary Table 1:** Diet composition and calorie content for diets used in this study.

**Supplementary Table 2:** Number of mice for each study cohort.

**Supplementary Table 3:** Number of mice for each figure.

**Supplementary Table 4:** Primer sequences for each gene measured by qPCR.

**Supplementary Table 5:** Hepatic Log^2^ fold-changes and related p-values for comparisons of interest from transcriptomic analysis.

**Supplementary Table 6:** Hepatic Log_2_ fold-changes and related p-values for comparisons of interest from metabolomic analysis.

